# A novel *kit* mutant rat enables hematopoietic stem cell engraftment without irradiation

**DOI:** 10.1101/2023.09.12.556153

**Authors:** Ryuya Iida, Saeko Ishida, Jinxi Wang, Kosuke Hattori, Kazuto Yoshimi, Satoshi Yamazaki, Tomoji Mashimo

**Author notes:** These authors contributed equally (Ryuya Iida, Saeko Ishida).

## Abstract

Hematopoietic stem cell (HSC) transplantation is extensively studied in mouse models, but their limited scale presents challenges for effective engraftment and comprehensive evaluations. Rats, due to their larger size and anatomical similarity to humans, offer a promising alternative. In this study, we establish a rat model with the *Kit*^V834M^ mutation, mirroring *Kit*^W41^ mice often used in KIT signaling and HSC research. *Kit*^V834M^ rats are viable and fertile, displaying anemia and mast cell depletion similar to *Kit*^W41^ mice. The mutation affects myeloid cell proliferation and differentiation, as seen in the colony-forming unit granulocyte-macrophage assay. Importantly, *Kit*^V834M^ rats support donor rat-HSC engraftment without irradiation. Competitive transplantation assays reveal reduced reconstruction capacity in *Kit*^V834M^ HSCs. Leveraging the larger scale of this rat model enhances our understanding of HSC biology and transplantation dynamics, potentially advancing our knowledge in this field.

## Introduction

Hematopoietic stem cell (HSC) engraftment, or bone marrow cell (BMC) transplantation, is a clinical procedure utilized to address various hematological disorders and certain cancers. Animal models play an essential role in unraveling the complexities of hematopoiesis, refining transplantation protocols, and advancing therapeutic strategies. Although mice have emerged as the predominant model for HSC transplantation studies owing to their well-established genetic resources, availability of immunodeficient strains for xenotransplantation investigations, and extensive understanding of mouse hematopoiesis^1^, their small size poses limitations on the number of transplanted HSCs in a single recipient. This constraint can affect engraftment efficiency and, consequently, the overall success of the transplantation procedure. Moreover, this limitation restricts the amount of biological samples available for subsequent testing and evaluation. Given that rats are approximately ten times larger than mice, they represent suitable candidates for specific experimental applications, particularly in transplantation studies^2,3^. Furthermore, the anatomical similarities between rats and humans, notably in the areas of bone marrow (BM) and the immune system^4,5^, suggest a more comparable environment for the growth and development of HSCs. Although non-human primate models, such as macaques, offer a closer representation of human physiology and the human immune system, their application is constrained by ethical considerations and elevated costs^6^.

In HSC engraftment, KIT signaling facilitates the successful establishment of transplanted HSCs within the recipient’s BM niche. The receptor tyrosine kinase KIT, also referred to as c-Kit or CD117, is a transmembrane protein characterized by distinct domains: the extracellular domain, transmembrane domain, juxtamembrane (JM) domain, and intracellular tyrosine kinase domain (depicted in Figure 1A). Stem cell factor (SCF), in its role as a ligand for KIT, facilitates receptor dimerization upon binding. This activation initiates a sequence of events involving autophosphorylation and transphosphorylation of tyrosine residues in the activation loop (AL) and JM domain, consequently activating the tyrosine kinase and triggering downstream signaling pathways. Activation of KIT signaling in HSCs enhances their ability to remain in the BM environment and aids in their interaction with supportive stromal cells, which are crucial for the survival of transplanted HSCs^8^. This signaling also plays a role in migration and homing of HSCs from the BM to the peripheral blood^9^. Importantly, KIT signaling is essential for regulation of the self-renewal capacity of HSCs and their long-term repopulation in HSC transplantation^10,11^.

**Figure 1.**
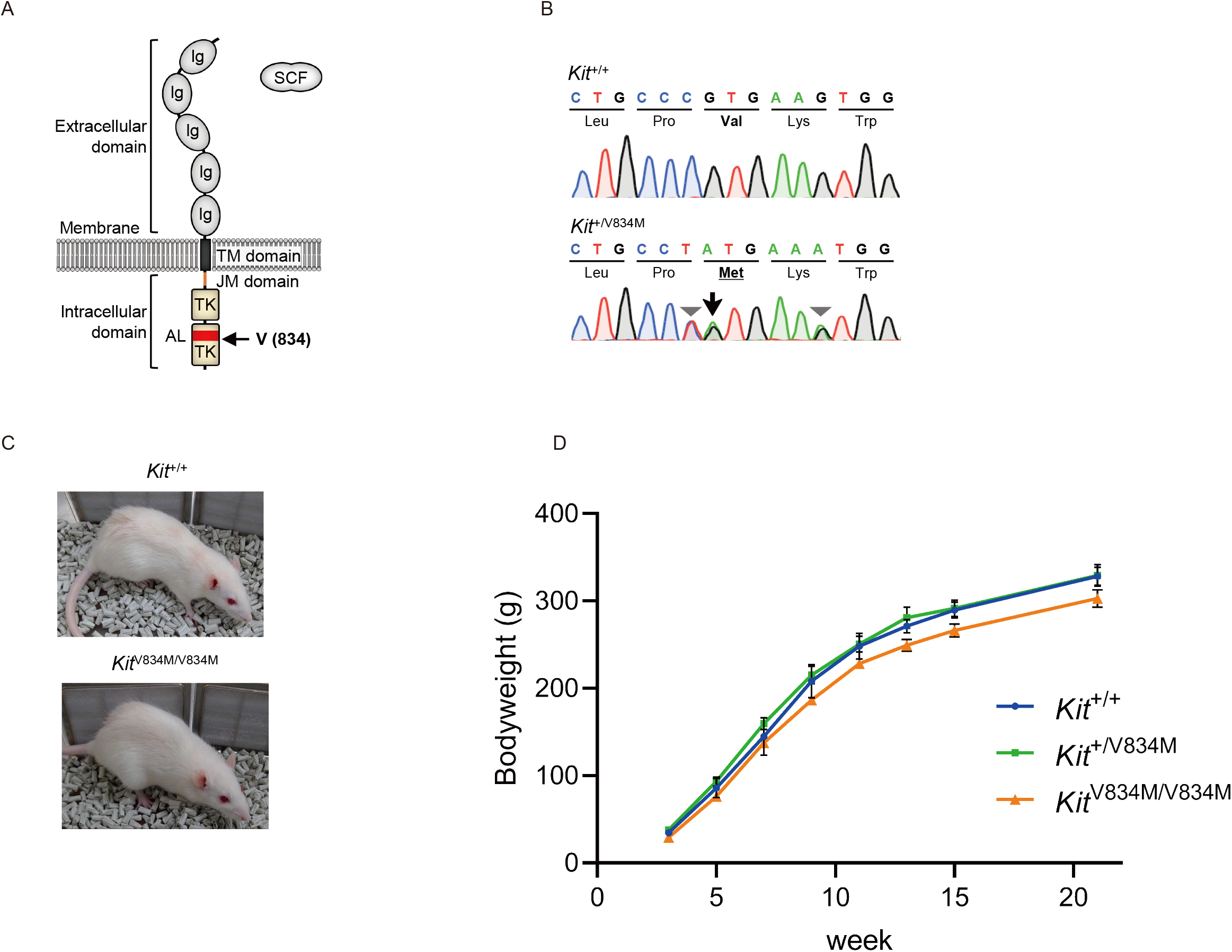
Generation of *Kit* ^V834M^ rats. (A)Structure of *Kit* and location of V834 in the activation loop (AL) region. The extracellular domain of *Kit* includes immunoglobulin-like (Ig) domains that bind ligands such as stem cell factor (SCF). The intracellular domain contains tyrosine kinase (TK) domains. AL spans into the C-lobe of the TK domain and maintains the TK in an inhibitory state with the juxtamembrane (JM) domain.(B)Sequence chromatograms of targeted region in *Kit*^+/+^ and *Kit*^+/V834M^ rats. Arrow indicates a missense mutation, c.2500G>A (V834M), and arrow heads indicate silent mutations, c.2499C>T and c.2505G>A. (C)Appearances of female *Kit*^+/+^ and *Kit*^V834M/V834M^ rats at 24 weeks of age. (D)Body weights of male *Kit*^+/+^ rats (n=3), *Kit*^+/V834M^ rats (n=6), and *Kit*^V834M/V834M^ rats (n=5). Graph error bars represent the mean⍰± ⍰SEM. Multiple comparisons of each group (*Kit*^V834M/V834M^ or *Kit*^+/V834M^ rats) versus *Kit*^+/+^ rats were conducted using Dunnett’s test. No significant differences were observed.

*Kit* mutant mouse models^12,13^ have been indispensable in unraveling the role of KIT in HSC transplantation. The *Kit* gene is located on the *White-Spotting (W)* locus in mice. Over 10 mouse variants with mutations in this locus have been documented, and the severity of their altered characteristics varies based on mutation type and genetic background^14–18^. Among these, *Kit*^W41^ mice have been extensively studied for the elucidation of KIT functions. *Kit*^W41^ mice carry a missense mutation (V845M) in the AL region, resulting in mild anemia that reflects HSC function^11,14,16,19^. These mice exhibit a nearly 50% reduction in the long-term HSC population compared with wild-type mice, and the self-renewing ability of their HSCs gradually decreases over time^11^. Notably, *Kit*^W41^ mice offer a supportive environment for HSC engraftment, eliminating the need for irradiation to repopulate the recipient’s hematopoietic system^13^. These distinctive traits render *Kit*^W41^ mice suitable for the investigation of HSC transplantation dynamics. In this study, to comprehensively understand KIT functions across different species and to evaluate the suitability of *Kit* mutant rats for HSC engraftment investigations, we established a novel *Kit* mutant rat model, denoted as F344-*Kit*^V834M^, which harbors an amino acid mutation that is identical to that of the well-studied *Kit*^W41^ mouse model.

## Results

### Generation of *Kit*^V834M^ rats

The KIT amino acid sequence is highly conserved among species. In the *Kit*^W41^ mouse, a substitution of valine to methionine occurs at position 835 (ENSMUST00000005815.7), which is equivalent to position 834 in rats (ENSRNOT00060032350.1). Valine at position 834 is located in the AL in the intracellular tyrosine kinase domain and plays important roles in the activation/deactivation processes, along with the JM domain^20^ (Figures 1A). SIFT analysis (http://sift.bii.a-star.edu.sg/), an evaluation method used to predict the impact of amino acid substitutions on protein function based on sequence homology and physical properties, predicts that the substitution at position 834 from valine to methionine is likely to affect protein function, with a SIFT score of 0.00.

To generate the *Kit*^V834M^ rat model, we used the CRISPR-Cas9 genome editing system. Our strategy involved designing a single-stranded oligodeoxynucleotide (ssODN) that introduced a missense mutation, along with two silent mutations in the seed sequence to prevent Cas9 from re-cutting the target sequence. We performed electroporation of the ssODN, crRNA, trans-activating crRNA, and Cas9 protein into 99 intact pronuclear-stage embryos obtained from inbred F344/NSlc strain rats, following a previously reported protocol^21,22^. Ninety-eight embryos survived the procedure, and we transferred these embryos into the oviducts of four pseudopregnant females. Subsequent Sanger sequencing analysis of the targeted region confirmed that three of the delivered pups carried the intended mutations. Germ line transmission of the mutations was further confirmed in the N1 generation of rats (Figure 1B).

### Normal fertility and weight gain observed in *Kit*^V834M^ rats

KIT is expressed in various stages of gametogenesis and is found in primordial germ cells, spermatogonia, and primordial and growing oocytes, indicating its involvement in different aspects of gamete development^23^. *Kit*^W41^ females exhibit significantly smaller and less active ovaries compared with wild-type controls^14^. In the case of the *Kit*^V834M^ rats generated here, fertility was observed and no abnormal deliveries were noted. Through interbreeding of *Kit*^+/V834M^ rats, we obtained offspring in the expected Mendelian ratio of wild-type (*Kit*^+/+^), heterozygous (*Kit*^+/V834M^), and homozygous (*Kit*^V834M/V834M^) rats (*Kit*^+/+^, n=23; *Kit*^+/V834M^, n=62; *Kit*^V834M/V834M^, n=37; P=0.4334; Chi-squared test). *Kit*^V834M/V834M^ rats also displayed fertility, and no differences in the size and histology of the testes and ovaries were observed compared with *Kit*^+/+^ rats (Figure S1A and S1B). Additionally, there were no differences in appearance between *Kit*^+/+^ and *Kit*^V834M/V834M^ rats (Figure 1C). Moreover, both male and female *Kit*^V834M/V834M^ rats exhibited similar body weights compared with those of control *Kit*^+/+^ rats (Figure 1D and S1C).

### *Kit*^V834M^ rats display age-dependent improvement in anemia and reduced mast cell count

HSCs are known for their capacity to differentiate into a spectrum of blood cell types, including red blood cells (RBCs) and immune cells such as mast cells, B cells, and T cells. KIT plays an indispensable role in HSC function, as substantiated by the mild anemia observed in *Kit*^W41^ mice^14,16^. Therefore, we evaluated critical hematological parameters—red blood cell count (RBC), hematocrit percentage (Ht or packed red cell volume), and mean cell volume (MCV)—in *Kit*^+/+^, *Kit*^+/V834M^, and *Kit*^V834M/V834M^ rats over time. Among male *Kit* ^V834M /V834M^ rats, a tendency toward mild macrocytic anemia was observed, characterized by reduced RBC scores and elevated MCV values—reminiscent of the pattern observed in *Kit*^W41^ mice. This anemic trend exhibited an inclination toward amelioration with increasing age (Figure 2A). A similar tendency was observed in female rats (Figure S2). Furthermore, we quantified the population of mast cells in the skin during postnatal weeks 8–9, as previously reported^24^. Male *Kit*^V834M/V834M^ rats displayed a marked reduction in the number of mast cells compared with *Kit*^+/+^ rats (Figure 2B). These findings collectively indicate potential functional deficiencies in the HSCs of *Kit*^V834M^ rats. Notably, we found that the frequencies of CD11b^+^ (myeloid cells), CD3^+^ (T cells), and CD45RA^+^ (B cells) among peripheral blood leukocytes, represented as a percentage of total CD45^+^ leukocytes, remained comparable between *Kit*^+/+^ and *Kit*^V834M/V834M^ rats (Figure 2C).

**Figure 2.**
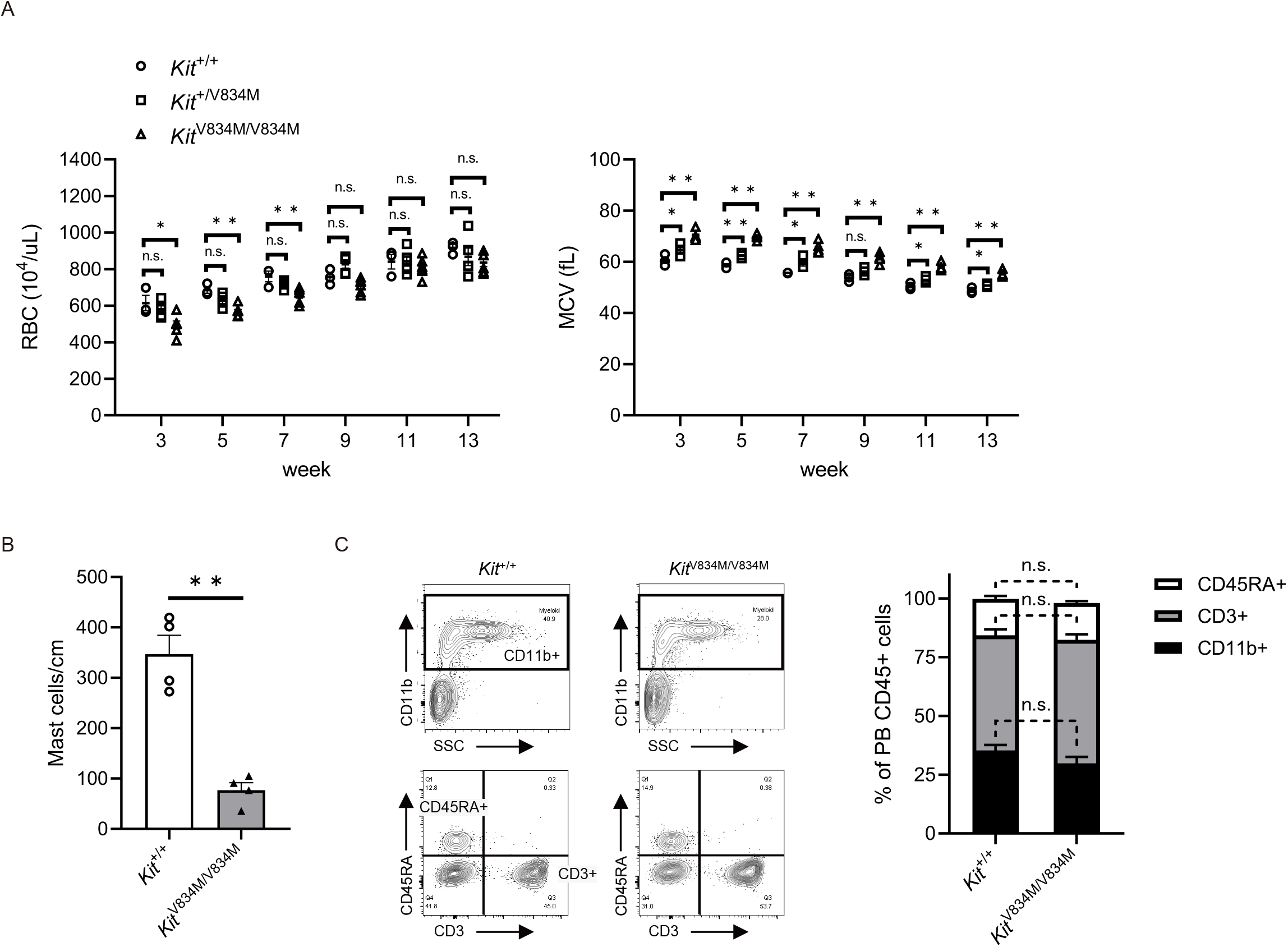
Anemia and decreased mast cell number in *Kit* ^V834M^ rats. (A)Hematological parameters of male *Kit*^+/+^ rats (n=3), *Kit*^+/V834M^ rats (n=5), and *Kit*^V834M/V834M^ rats (n=6). Graphs show mean cell volume (MCV, left) and red blood cell count (RBC, right). Multiple comparisons of each group (*Kit*^V834M/V834M^ or *Kit*^+/V834M^ rats) versus *Kit*^+/+^ rats were conducted using Dunnett’s test. Graph error bars represent the mean ⍰± ⍰SEM. *P<0.05; **P<0.01; n.s.: not significant.(B)The number of mast cells in the dorsal skin of 8- to 9-week-old male *Kit*^+/+^ rats (n=4) and *Kit*^V834M/V834M^ rats (n=4). Graph error bars represent the mean ⍰± ⍰SEM. **P<0.01, Student’s t- test.(C)Left: representative flow cytometry plots of peripheral blood from 20- to 25-week-old *Kit*^+/+^ and *Kit*^V834M/V834M^ rats. Right: frequencies of CD11b^+^ (myeloid cells), CD3^+^ (T cells), and CD45RA^+^ (B cells) peripheral blood leukocytes as percentages of all CD45^+^ leukocytes (n=4 for each). Graph error bars represent the mean ⍰± ⍰SEM. n.s.: not significant, Student’s t-test.

### Diminished differentiation potential observed in *Kit*^V834M^ HSCs

To comprehensively evaluate the functional competence of *Kit*^V834M/V834M^ (*Kit*^V834M^) HSCs, we conducted a colony-forming unit granulocyte-macrophage (CFU-GM) assay. In this assay, BM cells derived from *Kit*^V834M^ and wild-type *Kit*^+/+^ (WT) rats were cultured in semi-solid media with SCF, interleukin 3 (IL-3), and granulocyte-macrophage colony-stimulating factor (GM-CSF). After nine days of culture, the number of high proliferative potential (HPP) colonies, encompassing those exceeding 1 mm in diameter, was counted in accordance with established procedures^25^ (Figure 3A). Comparing the two groups, the total number of colonies appeared consistent between *Kit*^V834M^ and WT rats (Figure 3B). However, a notable difference emerged regarding HPP colonies: *Kit*^V834M^ rats exhibited a significantly diminished number of HPP colonies in contrast to WT rats (Figure 3C). These findings imply that the functional aberrations observed in *Kit*^V834M^ HSCs are not exhibited in their self-renewal or proliferation capabilities, but rather in their capacity to differentiate into diverse myeloid cell lineages.

**Figure 3.**
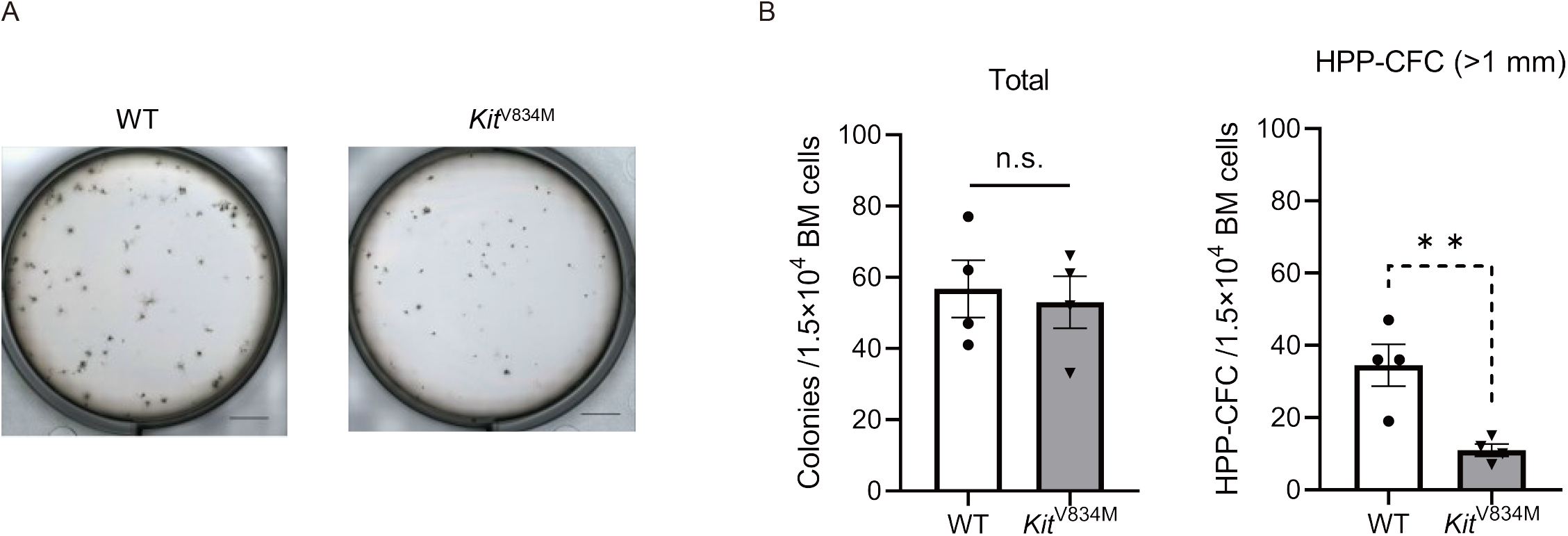
Lower differentiation potential observed in *Kit* ^V834M^HSCs. (A) Representative photographs of CFU-GM colonies at 9 days after seeding of bone marrow cells (BMCs) from WT (n=4) and *Kit*^V834M^ (n=4) rats. Bar graphs showing numbers of (B) total colonies and (C) HPP colonies with diameters greater than 1 mm. Scale bar indicates 5 mm. Graph error bars represent the mean ⍰± ⍰SEM. **P<0.01; n.s.: not significant, Student’s t-test.

### Syngeneic engraftment of donor BM Cells in *Kit*^V834M^ rats without irradiation

Considering the capacity of *Kit*^W41^ mice to foster engraftment of murine HSCs without the need for host irradiation^26^, we evaluated the potential of *Kit*^V834M^ rats as recipients for syngeneic BM cell transplantation. We introduced 2×10^7^ BM cells derived from F344-*Rosa26*^EGFP^ rats into the tail vein of *Kit*^V834M^ and WT rats without irradiation (Figure 4A). A longitudinal analysis of peripheral blood chimerism revealed a significant elevation in the average proportion of EGFP-positive cells among CD45^+^ peripheral blood cells, representing nucleated HSCs in *Kit*^V834M^ rats. This elevation reached 37.3 % by the 16th week post-transplantation. In contrast, EGFP^+^ donor cells were rarely observed in WT rats (Figure 4B). To characterize EGFP^+^ nucleated HSCs, we examined the expression of CD11b, CD3, and CD45RA, which are recognized markers for myeloid cells, T cells, and B cells, respectively. EGFP^+^ chimerism was exhibited in progressive increments within each of the CD11b^+^, CD3^+^, and CD45RA^+^ cell populations in *Kit*^V834M^ rats (Figure 4B). We further evaluated chimerism in the BM, spleen, and thymus. In the BM of *Kit*^V834M^ rats, the mean proportion of EGFP^+^CD45^+^ cells was 51.1 %, whereas these cells were rarely observed in WT rats (Figure 4C). Additionally, EGFP^+^CD45^+^ cells were observed in the spleen and thymus of *Kit*^V834M^ rats, but were rarely observed in the spleen and thymus of WT rats (Figures S3B).

**Figure 4.**
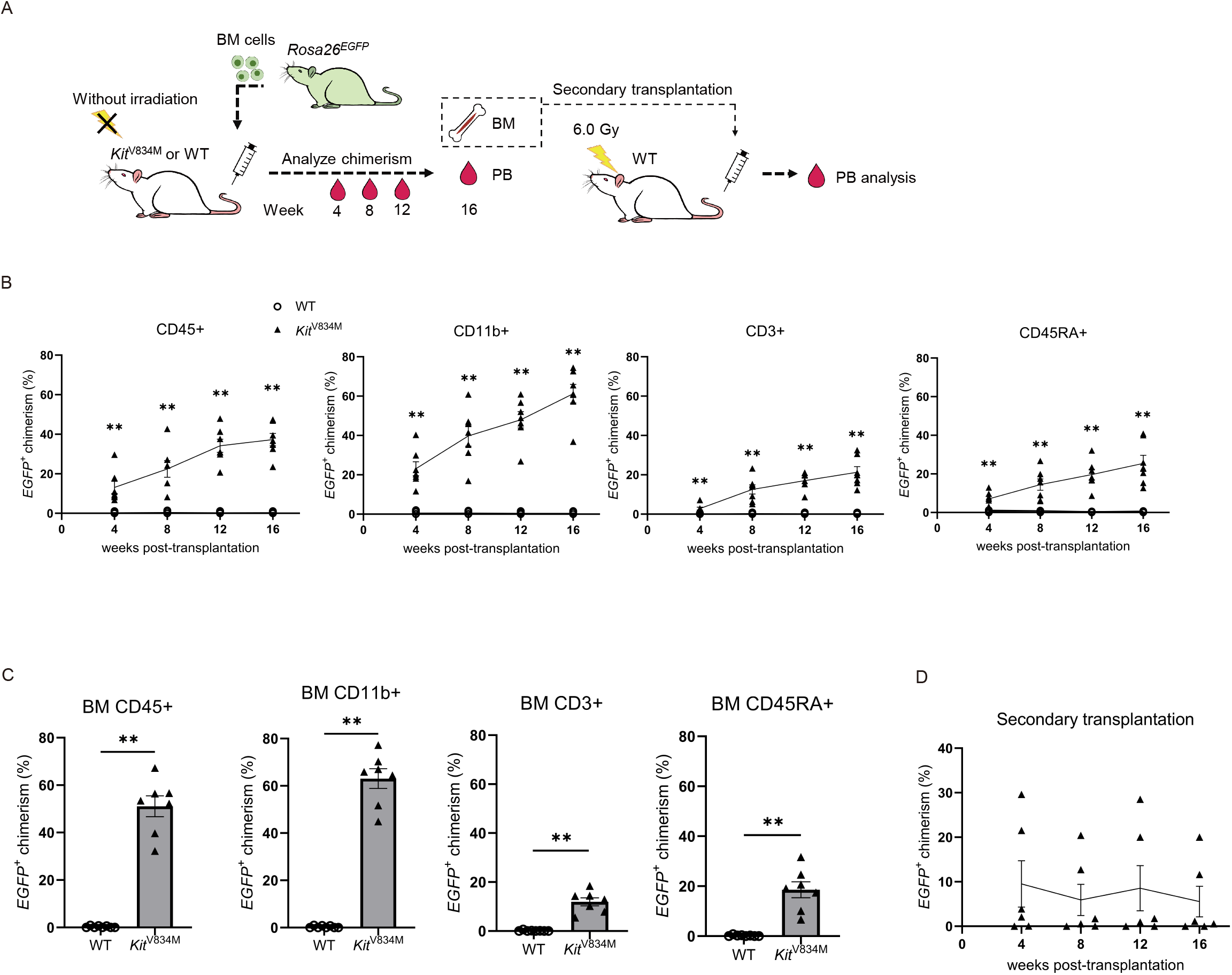
Successful syngeneic engraftment of donor BM Cells in *Kit*^V834M^ rats without irradiation. (A)Overview of syngeneic BM transplantation protocol.(B)Percentages of donor-derived EGFP^+^ cells in peripheral blood CD45^+^ (leukocytes), CD11b^+^ (myeloid cells), CD3^+^ (T cells), and CD45RA^+^ (B cells) cells of primary transplant recipients.(C)Percentages of donor-derived EGFP^+^ cells in BM of primary transplant recipients at 16 weeks post-transplantation.(D)Percentages of donor-derived EGFP^+^ cells in peripheral blood CD45^+^ cells of secondary transplant recipients. Graph error bars represent the mean ⍰± ⍰SEM. *P<0.05; **P<0.01; n.s.: not significant, Student’s t-test.

To evaluate the self-renewal capacity of engrafted HSCs in *Kit*^V834M^ rats, we harvested BM cells from *Kit*^V834M^ rats that had previously received EGFP^+^ BM cells. These harvested BM cells were subsequently re-transplanted into irradiated F344 rats. Following transplantation, EGFP^+^CD45^+^ cells emerged in the peripheral blood of the host F344 rats (Figure 4D). This observation indicates that the donor F344-*Rosa*26^EGFP^ HSCs within *Kit*^V834M^ rats had undergone self-renewal within the BM, thereby generating HSCs capable of repopulating secondary hosts.

### Diminished competitive repopulating capacity of *Kit*^V834M^ HSCs

To test our hypothesis that *Kit*^V834M^ HSCs exhibit a reduced capacity for competitive repopulation in comparison with WT HSCs, we conducted a competitive transplantation assay. Sublethally irradiated female WT rats were subjected to transplantation, receiving a 1:1 ratio of male *Kit*^V834M^ and F344-*Rosa*26^EGFP^ BM cells (Figure 5A). At 20 weeks post-transplantation, peripheral blood and BM cells were collected. Flow cytometry analysis showed that over 50% of CD45^+^ cells in both peripheral blood and BM samples were GFP^+^ cells (Figure 5B). In order to evaluate the chimerism characterizing each donor cell population, a quantitative real-time PCR assay was undertaken. Intriguingly, despite the transplantation of an identical BM cell quantity, the predominant cellular constituents retrieved from peripheral blood were derived from F344-*Rosa*26^EGFP^ rats (Figure 5C).

**Figure 5.**
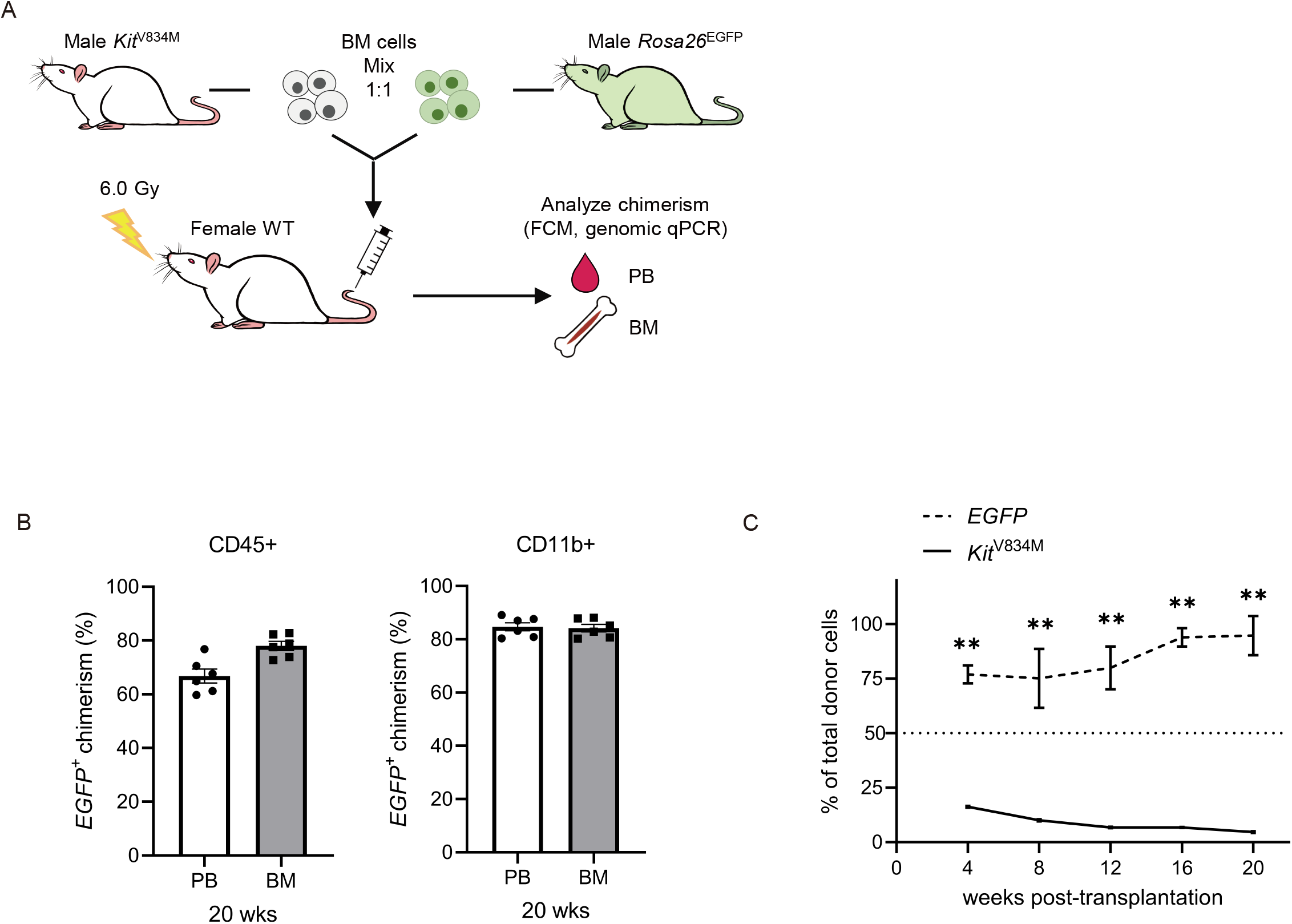
Reduced competitive repopulating ability of *Kit* ^V834M^ HSCs. (A) Overview of competitive transplantation assay protocol. (B) Percentages of *Rosa*26^EGFP^ in peripheral blood and BM CD45^+^, CD11b^+^ cells at 20 weeks post-transplantation measured by flow cytometry analysis. (C) Percentages of *Kit*^V834M^ and *Rosa*26^EGFP^ in donor-derived peripheral blood cells were measured at 4, 8, 12, 16, and 20 weeks after transplantation using genomic quantitative real-time PCR. Graph error bars represent the mean ⍰ ± ⍰ SEM. *P<0.05; **P<0.01; n.s.: not significant, Student’s t-test.

## Discussion

Rats and mice, the representative experimental animals for laboratory research, extend distinct advantages in diverse scientific fields. The larger size of rats, approximately tenfold that of mice, affords them special relevance in fields such as toxicology, physiology, and transplantation. Notably, the larger size facilitates vein accessibility for blood sampling, a convenience that eludes mouse research. In HSC engraftment, irradiation has traditionally been employed for myeloablative conditioning to create a receptive space within the recipient’s BM. Nonetheless, the collateral damage of irradiation includes cell necrosis, apoptosis, and the potential for complications such as infections. To address these concerns, we generated *Kit*^V834M^ rats, which offer an alternative approach for HSC transplantation that removes the need for irradiation. Notably, this approach finds parallel ground in *Kit*^W41^ mouse models harboring the identical *Kit*^V834M^ mutation, underscoring the robust potential to facilitate HSC transplantation without resorting to irradiation^13^.

Both the *Kit*^V834M^ rats and *Kit*^W41^ mice exhibited similar characteristics with respect to viability, fertility, and anemia, as shown in Figure 1. However, we also noticed some differences. One noticeable difference was in the reproductive organs. *Kit*^W41^ mice had smaller ovaries and fewer developing follicles, characteristics that were not observed in *Kit*^V834M^ rats. This observation might indicate that the effects of the *Kit* mutation on reproductive development could vary between species. Both *Kit*^V834M^ rats and *Kit*^W41^ mice had macrocytic anemia, but the anemia improved as *Kit*^V834M^ rats got older. *Kit*^Ws^rats^24,27^, which have a 12-base deletion in the AL region, have a diluted coat color and a large white spot on the belly, as seen in *Kit*^W^ and *Kit*^Wv^ mice. In the BN genetic background, homozygous mutation at the *Kit*^Ws^ mutant site seems to be fatal in rats; however, this is not the case in the Donryu genetic background. Unlike *Kit*^W^ and *Kit*^Wv^ mice, *Kit*^Ws^ rats exhibit a gradual improvement in anemia^24^, similar to our *Kit*^V834M^ rats. The colony-forming unit-erythroid (CFU-E) and/or its immediate precursors in *Kit*^Ws^ rats are more sensitive to erythropoietin than those of *Kit*^Wv^ mice^24^. Thus, there might be differences in erythropoiesis between mice and rats. These findings highlight the importance of studying different gene mutations, genetic backgrounds, and species to comprehensively understand the effect of *Kit* mutations.

HSCs give rise to progenitor cells, including common myeloid progenitors (CMP) and common lymphoid progenitors, through the multipotent progenitor cell stage. In our experiment involving F344-*Rosa*26^EGFP^ BM transplantation into recipient *Kit*^V834M^ rats, we examined the chimerism of donor EGFP^+^ cells in various cell populations (Figure 4). Interestingly, the chimerism of donor EGFP^+^ cells was lower in CD3^+^ (T cell) and CD45RA^+^ (B cell) populations compared with the CD11b^+^ (myeloid cells) population in both peripheral blood and BM (Figure 4B and C). These findings suggest a potential impairment in the generation or expansion of myeloid cells derived from CMP with the *Kit*^V834M^ mutation, which is consistent with previous reports in *Kit* mutant mice^28^. The CFU-GM assay further illustrated that *Kit*^V834M^ HSCs maintain the ability to differentiate into CMP, albeit with a slower growth rate (Figure 3C). Despite their capacity to differentiate, the slower proliferation of mutant HSCs hampers their complete repopulation of the recipient’s BM cells. In the competitive transplantation assay, when both donor *Kit*^V834M^ and *Rosa*26^EGFP^ HSCs were transplanted into recipient animals, only a small proportion of *Kit*^V834M^ cells were detected in the peripheral blood of recipients via qPCR (Figure 5C). The limited chimerism of *Kit*^V834M^ cells in recipients indicates reduced engraftment efficiency or a competitive disadvantage compared with wild-type HSCs. Notably, in the case of *Kit*^W41^ mice, successful reconstitution was achieved in lethally irradiated wild-type mice by transplanting a higher ratio of *Kit*^W41^ to wild-type BM cells (e.g., 50:1 ratio). However, the chimerism decreased with age, particularly in myeloid cells, which suggests the reduced reconstitution ability of *Kit*^W41^ HSCs in serially transplanted recipients^28^. These findings highlight the importance of studying the engraftment abilities of *Kit* mutant HSCs, and enhance our understanding of the functional implications of the *Kit*^V834M^ mutation on HSC behavior and transplantation outcomes.

The *Kit*^W41^ mutation has been integrated with NOD-Prkdc, Il2rg (NSG) mice to create a novel model for HSC xenotransplantation^13^. This immunodeficient mouse model enables the engraftment of human HSCs in the mouse BM without the need for irradiation. This provides an invaluable tool for researchers studying HSC xenotransplantation. Beyond mice, rats with severe combined immunodeficiency (SCID) with Il2rg and/or Rag2 gene knockouts^29^ have also been utilized in various research fields, such as the engraftment of cancer cells derived from patients^30^, as well as for the successful transplantation of human induced pluripotent stem cell-derived airway epithelial cell sheets^31^, miniature human livers^32^, and human mesenchymal progenitor cells within humanized bone constructs^33^. By combining the *Kit*^V834M^ mutation with SCID rats, we can anticipate further advancements in the generation of humanized rat models for HSC research.

In conclusion, we demonstrate a novel *Kit* mutant rat, *Kit*^V834M^, which is partially distinct from previously reported *Kit* mutant mice and rats. While *Kit*^V834M^ rats show anemia and their HSCs are less competitive, these rats are viable and fertile, even in a homozygous state. We also found that *Kit*^V834M^ rats permit allogenic engraftment of HSCs in the absence of irradiation. *Kit*^V834M^ rats will provide new understanding of *Kit* function and experimental tools for various research fields, including hematology, immunology, and translational research.

## Acknowledge

We thank Junko Imamura, Yuko Yamauchi, and Eri Ezawa for expert technical support in rat maintenance and genotyping. We also thank Tamami Denda and Dr. Toshihiro Kobayashi for help with histological analysis. We acknowledge the IMSUT FACS Core laboratory for assistance with flow cytometry analysis.

## Author Contributions

RI, SI, SY and TM designed experiments. RI and SI conducted the experiments. JW, KH and KY generated rats. RI, SI, JW, KH, KY, SY and TM contributed to the writing of the paper.

## Figure legends

**Figure S1. Normal gonadal histology and weight gain observed in *Kit*** ^**V834M**^**rats**

Hematoxylin-and eosin (HE) stained paraffin-embedded sections of ovaries (A) ovaries and (B) testes from *Kit*^V834M^ and WT rats. (C) Body weights of female *Kit*^+/+^ rats (n=5), *Kit*^+/V834M^ rats (n=8), and *Kit*^V834M/V834M^ rats (n=3). Graph error bars represent the mean ⍰ ± ⍰ SEM. Multiple comparisons of each group (*Kit*^V834M/V834M^ or *Kit*^+/V834M^ rats) versus *Kit*^+/+^ rats were conducted using Dunnett’s test. No significant differences were observed.

**Figure S2. Anemia observed in female *Kit*^V834M^ rats** Hematological parameters of female *Kit*^+/+^ rats (n=5), *Kit*^+/V834M^ rats (n=5), and *Kit*^V834M/V834M^ rats (n=3). Graphs show mean cell volume (MCV, left) and red blood cell count (RBC, right). Multiple comparisons of each group (*Kit*^V834M/V834M^ or *Kit*^+/V834M^ rats) versus *Kit*^+/+^ rats were conducted using Dunnett’s test. Graph error bars represent the mean ⍰ ± ⍰ SEM. *P<0.05; **P<0.01; n.s.: not significant.

**Figure S3. Successful engraftment observed in the spleen and thymus of *Kit*^V834M^ rats** EGFP^+^CD45^+^ cells were observed in the spleen and thymus in *Kit*^V834M^ rats, but not in WT rats

## STAR ⍰ Methods

### Key resources table

| REAGENT or RESOURCE | SOURCE | IDENTIFIER |
| --- | --- | --- |
| **Antibodies** |  |  |
| APC/Cyanine7 anti-rat CD45 Antibody (Clone OX-1) | Biolegend | Cat# 202216; RRID: AB_1236411 |
| PE Mouse Anti-Rat CD11b (Clone WT.5) | BD Bioscience | Cat# 562105; RRID: AB_10896966 |
| PE anti-rat CD3 Antibody (Clone 1F4) | Biolegend | Cat# 201412; RRID: AB_2563273 |
| PerCP/Cyanine5.5 anti-rat CD45RA Antibody (Clone OX-33) | Biolegend | Cat# 202318; RRID: AB_2565945 |
| **Chemicals, peptides, and recombinant proteins** |  |  |
| Alt-R™ S.p. Cas9 Nuclease V3 | Integrated DNA Technologies | 1081058 |
| Isoflurane | VIATRIS | N/A |
| 4%-Paraformaldehyde Phosphate Buffer Solution | NACAL TESQUE | 09154-85 |
| Sucrose | WAKO Fuji Film | 193-00025 |
| D-PBS(-) without Ca and Mg, liquid | NACAL TESQUE | 14249-24 |
| 10× RBC Lysis Buffer (Multi-species) | eBioscience | 00-4300-54 |
| PI solution | Dojindo | P378 |
| HiDi DNA polymerase | myPOLS Biotec Gmbh | 9001M |
| GreenDye 20× | myPOLS Biotec Gmbh | 2000M |
| MethoCult™ GF R3774 | Stem Cell technologies | 03774 |
| Maxwell® RSC Whole Blood DNA Kit | Promega | AS1520 |
| **Oligonucleotides** |  |  |
| gRNA and ssODN used for CRISPR/Cas9 mediated knock-in | Table S1 | N/A |
| Primers for genotyping | Table S2 | N/A |
| Primers for qPCR | Table S3 | N/A |
| **Software and algorithms** |  |  |
| FlowJo software 10.7.2. | Flowjo | N/A |
| GraphPad Prism version 9 | GraphPad Software | N/A |
| **Other** |  |  |
| NEPA21 Super Electroporator Type II | NEPA Gene | N/A |
| FUJI TUBE (EDTA-2K) | Fuji Film | N/A |
| MEK-6558 Celltacα | Nihon Kohden | N/A |
| Nanophotometer-N60 Touch | IMPLEN | N60-TOUCH |
| Maxwell RSC Instrument | Promega | AS4500 |
| CytoFLEX LX N3-V5-B3-Y5-R3-I0 Flow Cytometer | Beckman Coulter | C00446 |
| CFX Connect Real-Time PCR System | Bio-Rad Laboratories | N/A |

## Resource availability

### Lead contact

Further information and requests for resources and reagents should be directed to and will be fulfilled by the Lead Contact Tomoji Mashimo.

### Materials availability

This study utilized F344-*Kit*^V834M^ that our lab generated. This resource is available upon request to the lead contact as indicated above, and will be deposited at the National Bio Resource Project Rat in Japan (www.anim.med.kyoto-u.ac.jp/nbr).

## Experimental model and subject details

### Animals

All rats were housed and maintained in the animal facilities at the Institute of Medical Science, the University of Tokyo. Rats were kept on a 12 h light/dark cycle with food and water ad libitum under specific pathogen-free conditions. All animal care and experimental procedures were approved by the Institutional Animal Care and Use Committee of the University of Tokyo.

### Generation of mutant rats

To generate F344-*Kit*^em1larc^ rats, a guide RNA, single-stranded oligodeoxynucleotide (ssODN), and Cas9 protein (Alt-R S.p. Cas9 Nuclease V3, Integrated DNA Technologies) were electroporated into intact pronuclear-stage embryos of F344/Jcl rats^21,22^. The gRNA and ssODN sequences are shown in Table S1. All surviving embryos were transferred into the oviducts of pseudopregnant females. Genomic DNA of founder rats was extracted from ear biopsies. The gRNA target site was amplified using primers listed in Table S2. Successful germline transmission was confirmed by DNA sequencing of the first backcross progenies.

### Histological analysis

Rats were deeply anesthetized by isoflurane and then perfused transcardially with 4% paraformaldehyde (PFA). Testes and ovaries were taken from 10-week-old rats, and pieces of the dorsal skin were taken from 8- to 9-week-old male rats. The samples were post-fixed by 4% PFA and embedded in paraffin. Sections of testis and ovary (10 μm thick) were stained with hematoxylin and eosin. Skin sections (4 μm thick) were stained with alcian blue and Kernechtrot. Differentiation of germ cells in the testis and ovary was evaluated and the number of mast cells was counted using a microscope (BZ-800, Keyence) as previously reported^24^.

### Hematologic examination

Peripheral blood samples were obtained from tail veins every two weeks. The blood parameters, RBC, and MCV were determined by Celltacα (MEK-6558, Nihon Kohden).

### Colony-forming assay

For colony assays, 1.5×104 BM cells were seeded in duplicates with 1.1 ml Methocult medium (Methocult GF R3774, Stem Cell Technologies) into 6-well plates and further cultivated at 37 °C in a 5% CO2 atmosphere. The number of total colonies and HPP colony-forming cells, which generate a colony with a diameter >1 mm, were evaluated on day 9 of culture.

### Bone marrow transplantation

BM cells were isolated from the femur and tibia of rats, as previously reported^34^. For BM serial transplantation experiments, 2.0×10^7^ BM cells from male F344-*Rosa*26^EGFP^ rats were transplanted into 5-week-old male F344-*Kit*^V834M^ rats without irradiation. Chimerism of GFP^+^ cells in peripheral blood was monitored every 4 weeks after transplantation by flow cytometric analysis. Sixteen weeks after transplantation, primary recipient rats were sacrificed and chimerism in BM, spleen, and thymus cells was analyzed by flow cytometry. The cells were also transplanted into irradiated (6.0 Gy) secondary recipient F344/Jcl rats.

For the competitive BM transplantation assay, 1.0×107 BM cells were harvested from male F344-*Kit*^V834M^ rats and were transplanted together with 1.0×107 BM cells from male F344-*Rosa*26^EGFP^ into irradiated (6.0 Gy) recipient female F344/Jcl rats via tail veins. Chimerism in peripheral blood cells was monitored every 4 weeks by flow cytometric analysis and genomic quantitative real-time PCR.

### Flow cytometry analysis

Peripheral blood was collected from the tail vein. To remove RBCs, cells were treated with 1X RBC Lysis Buffer (eBioscience) according to the manufacturer’s instructions. Single-cell suspensions were prepared from BM, spleen, and thymus by standard procedures. For flow cytometric analysis, the following antibodies were used: APC/Cyanine7-CD45 (Biolegend, 202216), PE-CD11b (BD Biosciences, 562105), PE-CD3 (Biolegend, 201412), and PerCP/Cyanine5.5 CD45RA (Biolegend, 202318). Cells were stained with appropriate antibodies in 100 μl PBS for 30 min on ice. Flow cytometry was performed on the CytoFLEX LX (Beckman Coulter). Propidium iodide was used to exclude dead cells. The results were analyzed with FlowJo software 10.7.2.

### Quantitative real-time PCR

The chimerism of donor cells was evaluated by quantitative PCR. To distinguish male donor cells from female recipient cells, Y chromosome specific gene Zfy1 was used as previously reported^35^. HiDi (high single nucleotide discrimination) DNA polymerase (myPOLS Biotec Gmbh) was used to amplify *Kit*^V834M^ cells specifically. The used primers are listed in Table S3. The percentage of male donor-derived cells, *Kit*^V834M^ and F344-*Rosa*26^EGFP^, in the female recipient mice was calculated against standard curve with known percentage of male F344-*Kit*^V834M^ vs. male F344-*Rosa*26^EGFP^ DNA.

## Statistical analysis

Statistical details of each experiment can be found in each figure legend. Statistical analysis was performed using Graph Pad Prism 9 software. P<0.05 was considered statistically significant.

