## Supplementary figures and images for "A novel *kit* mutant rat enables hematopoietic stem cell engraftment without irradiation"

### Figure S

Figure S1

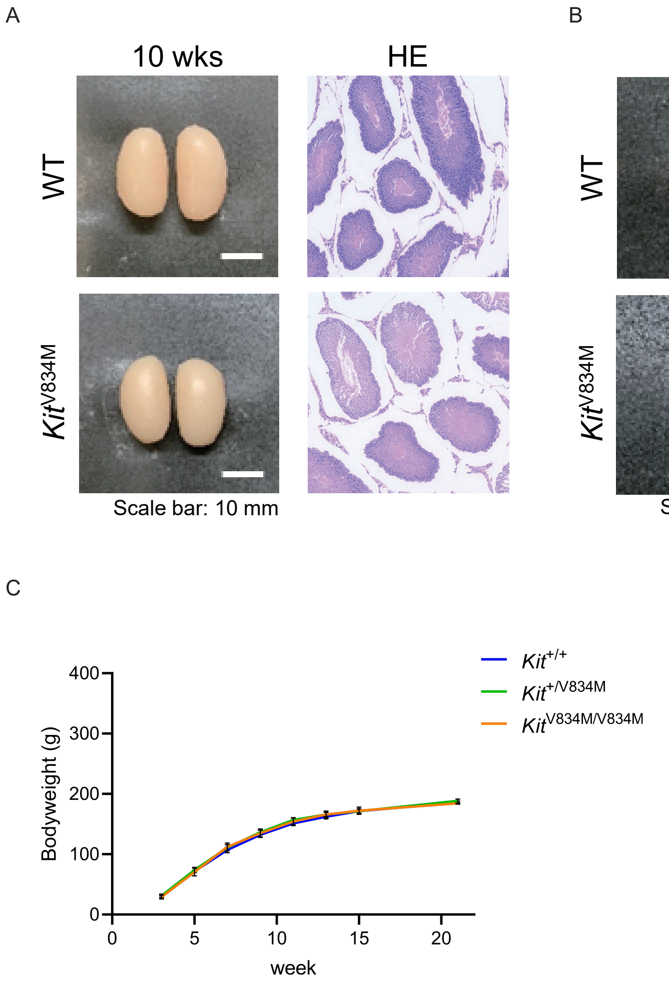


Figure S2

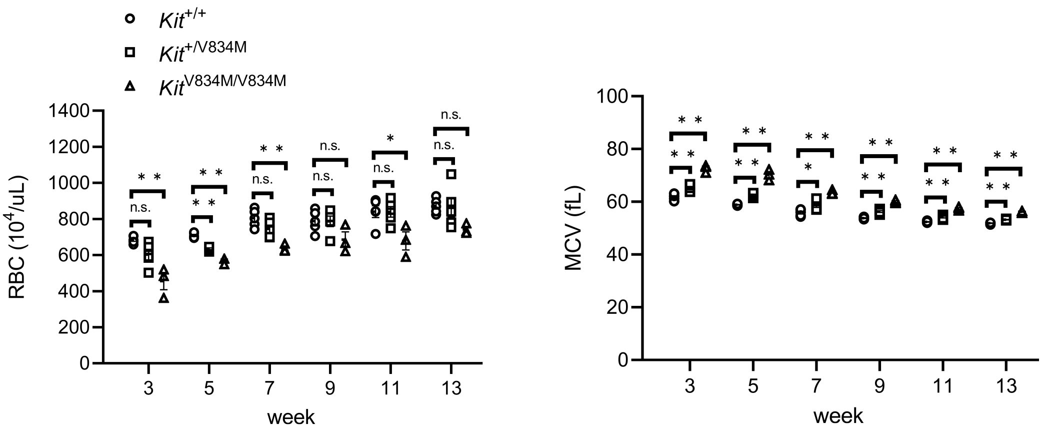


Figure S3

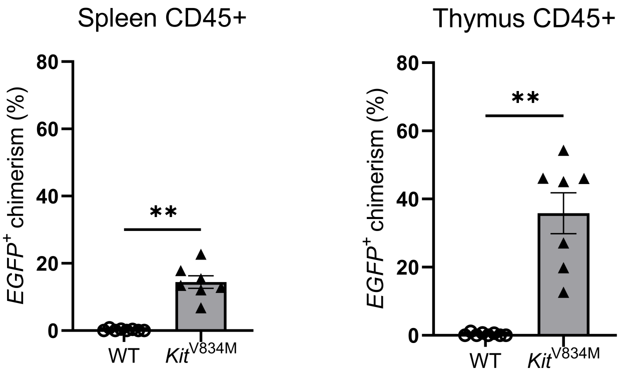
